# Neural representations of non-native speech reflect proficiency and interference from native language knowledge

**DOI:** 10.1101/2023.04.15.537014

**Authors:** Christian Brodbeck, Katerina Danae Kandylaki, Odette Scharenborg

## Abstract

Learning to process speech in a foreign language involves learning new representations for mapping the auditory signal to linguistic structure. Behavioral experiments suggest that even listeners that are highly proficient in a non-native language experience interference from representations of their native language. However, much of the evidence for such interference comes from tasks that may inadvertently increase the salience of native language competitors. Here we tested for neural evidence of proficiency and native language interference in a naturalistic story listening task. We studied electroencephalography responses of native Dutch listeners to an English short story, spoken by a native speaker of either American English or Dutch. We modeled brain responses with multivariate temporal response functions, using acoustic and language models. We found evidence for activation of Dutch language statistics when listening to English, but only when it was spoken with a Dutch accent. This suggests that a naturalistic, monolingual setting decreases the interference from native language representations, whereas an accent in the listeners’ own native language may increase native language interference, by increasing the salience of the native language and activating native language phonetic and lexical representations. Brain responses suggest that words from the native language compete with the foreign language in a single word recognition system, rather than being activated in a parallel lexicon. We further found that secondary acoustic representations of speech (after 200 ms latency) decreased with increasing proficiency. This may reflect improved acoustic-phonetic models in more proficient listeners.

## 1 Introduction

A plethora of behavioral studies has shown that non-native speech processing is slower and more error-prone than native speech processing, even in highly proficient listeners, and especially in adverse listening conditions (see Garcia Lecumberri et al., 2010; Scharenborg & van Os, 2019 for a review). One reason for this is the influence of the native language on non-native listening (Cutler, 2012; Garcia Lecumberri et al., 2010). Our knowledge of the sounds of our native language influences how we perceive non-native sounds, which increases the number of misperceptions of sounds by non-native listeners compared to native listeners (Garcia Lecumberri et al., 2010). This problem percolates upwards in the recognition process, leading to the spurious activation of non-target words from both the non-native language (Cutler et al., 2006; Karaminis et al., 2022; Scharenborg et al., 2018) and the native language (Hintz et al., 2022; Marian & Spivey, 2003; Spivey & Marian, 1999; Weber & Cutler, 2004), slowing down word recognition and decreasing word recognition accuracy for non-native listeners (Broersma & Cutler, 2008, 2011; Drijvers et al., 2019; Perdomo & Kaan, 2021).

In native language processing, listeners have been found to employ predictive language models of speech at different levels in parallel, including the sublexical, word-form and sentence levels (Brodbeck et al., 2022; Z. Xie et al., 2023). Listeners that have no knowledge of the target language do not exhibit comparable responses (Tezcan et al., 2022). We thus hypothesized that acquiring a new language involves developing such predictive models, and that those models exhibit evidence for interference from the native language at the sublexical and word recognition levels. At the level of word recognition, we contrast two different possible mechanisms of native language interference (see Figure 5). The standard view is that Dutch and English word forms directly compete for recognition in one shared lexicon (Brysbaert & Duyck, 2010; Dijkstra et al., 2019). Alternatively, words from the two languages could be activated in segregated lexical systems, and interference would then only occur at the level of behavioral output (e.g., eye movements in a visual world study).

Previous research on native language interference typically focused on behavioral experiments using carefully crafted stimuli. However, recent results suggest that tasks which increase meta- linguistic awareness also increase the influence of the native language on non-native speech perception (Freeman et al., 2021). This may have led to an overestimation of the effects of native language interference. Here we use a naturalistic listening paradigm and measure neural responses to speech unobtrusively with electroencephalography (EEG), while native speakers of Dutch listen to an English story. We included two recordings of the story, spoken with an American vs. a Dutch accent, manipulating salience of the native language.

Non-native listeners are known to benefit from a non-native accent that corresponds to their own language background (Bent & Bradlow, 2003), although this is not always the case (Gordon-Salant et al., 2019; Hayes-Harb et al., 2008). This heterogeneity may be due to an interaction with proficiency in the target language: non-native listeners with higher proficiency in the target language tend to show a higher accuracy for native accents, whereas lower proficiency non-native listeners may benefit more from the accent of their own native language (Pinet et al., 2011; X. Xie & Fowler, 2013). We measured proficiency to determine whether effects of speaker accent change as a function of proficiency.

We investigated four related questions: 1) Is there evidence for parallel predictive language models in non-native listeners? 2) Do brain responses to non-native speech exhibit evidence for interference from native language statistics? 3) Do these effects depend on the accent of the speaker? 4) Do the effects change as a function of language proficiency, and is the effect of accent modulated by proficiency? I.e., do highly proficient listeners benefit more from a native accent (American accented English), and low proficiency listeners from an accent of their own native language (Dutch accented English)?

## 2 Materials and Methods

In order to measure neural representations during naturalistic non-native story listening, we used the multivariate temporal response function (mTRF) framework (Brodbeck et al., 2021; Lalor & Foxe, 2010). Participants listened to an approximately 12 minute long English story twice, once spoken with an American English accent, and once with a Dutch accent, with the order counterbalanced across participants. Using 5-fold cross-validation, mTRFs were trained to predict the EEG responses to each story separately from multiple predictor variables, reflecting different acoustic and linguistic properties of the stimuli (see Figure 1 and below). Predictor variables for English closely followed previously reported research (Brodbeck et al., 2022). The influence of native language (Dutch) knowledge on neural representations was assessed by generating additional predictors from Dutch language statistics. To determine which neural representations change as a function of non-native proficiency, the predictive power of the different groups of predictors across listeners was correlated with behavioral tests measuring non-native language proficiency.

**Figure 1.**
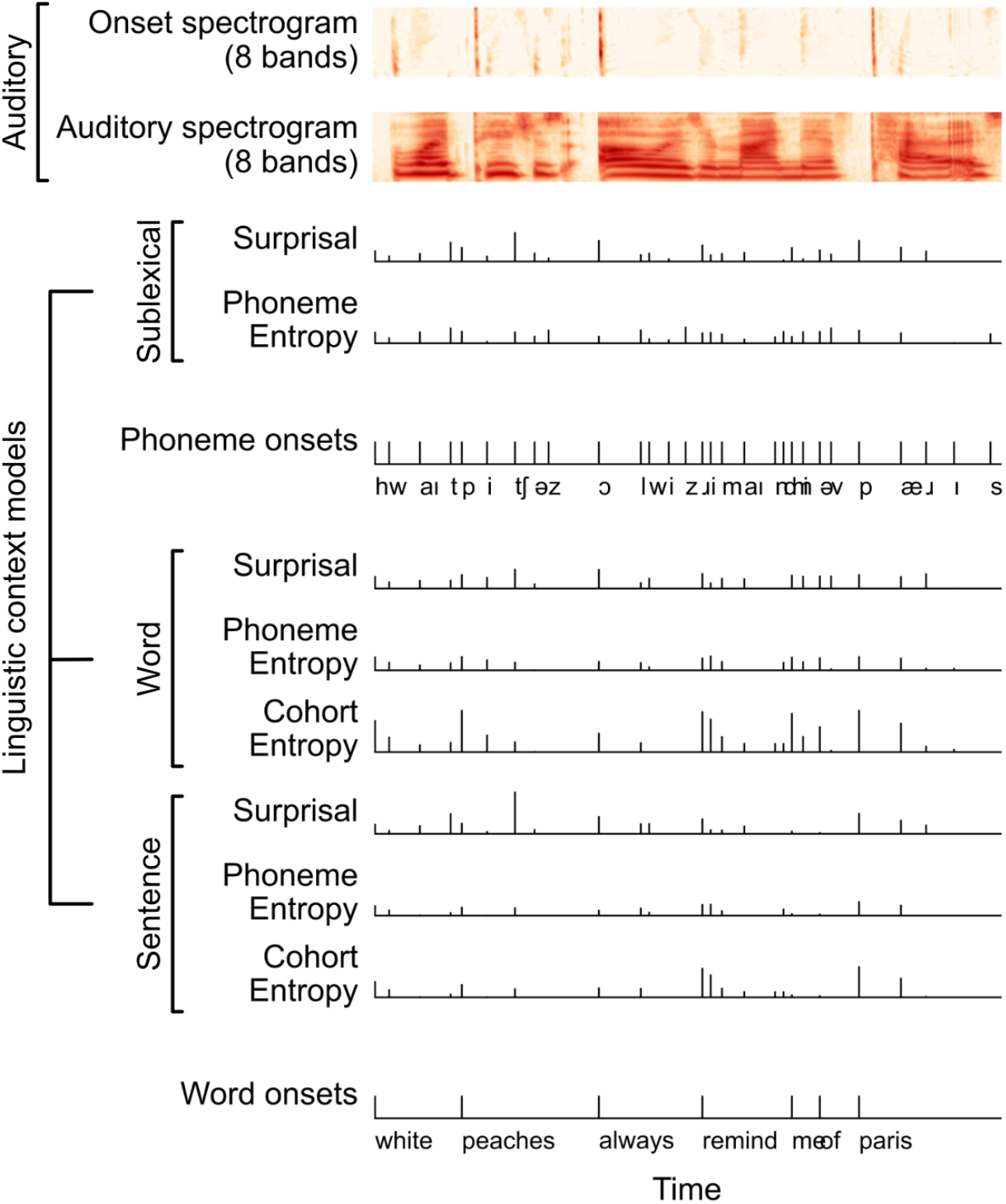
Analysis design: predictors and groups of predictors used to test specific hypotheses. Each predictor was constructed as a continuous time series, aligned with the stimuli and corresponding EEG responses. Both auditory predictors were reduced to 8 bands, equally spaced in equivalent rectangular bandwidth, to simplify the analysis computationally. Predictors were grouped into sets that reflect specific processes of interest, as indicated by brackets.

### 2.1 Participants

Forty-six Dutch non-native listeners of English from the Radboud University, Nijmegen, the Netherlands, subject pool participated in the experiment. Seven participants were excluded due to technical issues during data acquisition: for two participants, part of the EEG recordings was missing; for one participant the sound level was initially too low, so part of the English story was presented twice; for one participant the event codes were missing; for one participant the connection with the laptop was lost; for one participant the battery failed during the experiment; for one participant the behavioral data was missing. This left a sample of 39 participants (14 males, mean age: 21.6, standard deviation (SD): 2.7; range 18-29). The experiment consisted of two parts: a lexically-guided perceptual learning experiment followed by listening to the two stories. The lexically-guided perceptual learning experiment, which investigated the neural correlates underlying lexically-guided perceptual learning, was reported in Scharenborg et al. (2019). The participants reported here are a superset of those reported in Scharenborg et al. (2019). All participants were paid for participation in the experiment. No participants reported hearing or learning problems.

#### 2.1.1 Non-native proficiency: LexTale

General English proficiency of the Dutch non-native listeners of English was assessed using the standardized test of vocabulary knowledge, LexTale (Lemhöfer & Broersma, 2012). LexTale scores ranged from 46 (which corresponds to a “B1 and lower” level of proficiency according to the Common European Framework of Reference for Language) to 92 (which indicates a C1 and C2 level of proficiency or an “upper & lower advanced/proficient user”; note Lemhöfer and Broersma do not differentiate between C1 and C2 levels). The mean LexTale score was 73.3 (SD=11.0), which corresponds (roughly) to an upper intermediate or B2 level of proficiency (Lemhöfer & Broersma, 2012). All non-native English participants were taught English in high school for at least 6 years.

#### 2.1.2 Acoustic-phonetic aptitude: LLAMA_D

The LLAMA test (Meara et al., 2002) consists of five tests to assess aptitude for learning a foreign language, and is based on Carroll and Sapon (1959). The five tests assess different foreign language learning competencies, including vocabulary learning, grammatical inferencing, sound-symbol associations, and phonetic memory. Here we used the LLAMA_D sub-test, which assesses the ability to recognize auditory patterns, a skill that is essential for sound learning and ultimately word learning. We therefore refer to the LLAMA_D score as acoustic-phonetic aptitude.

During the test, participants first heard a list of words; in the second part of the test, participants heard new and repeated words, and were asked to indicate whether the stimulus was part of the initial target words. The words were synthesized using the AT&T Natural Voices (French) on the basis of flower names and natural objects in a Native American language of British Columbia, yielding sound sequences that are not recognizable as belonging to any major language family. The participants got feedback regarding the correctness of their answer after each trial. They scored points for correctly recognized target words and lost points for mistakes. This tested the ability to recognize repeated stretches of sound in an unknown phonology, which is an important skill for learning words in a foreign language (Service et al., 2022), and for distinguishing variants that may signal morphology (Rogers et al., 2017).

The LLAMA_D scores range between 0 and 100%, where 0-10 is considered a very poor score, 15-35 an average score (most people score within this range), 40-60 a good score, and 75-100 an outstandingly good score (few people manage to score in this range) (Meara, 2005). A previously reported average score is 29.3%, SD=11.4 (Rogers et al., 2017).

### 2.2 Stimuli

The short story was an excerpt from the book “Garlic and sapphires: The secret life of a critic in disguise” by Ruth Reichl (2005); specifically, chapter “The daily special”. We aimed to select a story on a neutral topic, while avoiding books that our participants would be familiar with. At the same time we wanted the story to be entertaining so that participants would be engaged with the story and would want to continue to listen.

The stories were read by a female native American speaker and a female Dutch speaker, both students at the Radboud University at the time of recording. Recordings were made in a sound-attenuated booth using a Sennheiser ME 64 microphone. Each speaker read the story twice. The story with the fewest mispronunciations was chosen for the experiment. Both stories were around 12 minutes long.

### 2.3 Procedure

Participants were tested individually in a sound-attenuated booth, comfortably seated in front of a computer screen. The two short stories were administered in a single session after the lexically- guided perceptual learning experiment reported previously (Scharenborg et al., 2019). The intensity level of both stories was set at 60 dB SPL and was identical for all participants. The stories were played with Presentation 17.0 (Neurobehavioral Systems, Inc.), and were presented binaurally through headphones.

Participants saw an instruction on the computer screen informing them that they would be listening to two short stories in English. To start the story, participants had to press a button. Once the story was finished, the participants were prompted to press another button to start the second story. The order of the presentation of the two stories was balanced across participants.

We recorded EEG activity continuously during the entire duration of the experiment from 32 active Ag/AgCI electrodes, placed according to the 10-10 system. The left mastoid was used as an online reference and the average of left and right mastoids as an offline reference. Eye movements were monitored with additional electrodes placed on the outer canthus of each eye and above and below the right eye. Impedances were generally kept below 5 KOhm. Data were sampled at 500 Hz after applying an online 0.016 – 125 Hz bandpass filter.

### 2.4 Analysis

#### 2.4.1 Preprocessing

EEG data were preprocessed with MNE-Python (Gramfort et al., 2014). Data were band-pass filtered between 1 and 20 Hz (zero-phase FIR filter with MNE-Python default settings), and biological artifacts were removed with Extended Infomax Independent Component Analysis (Bell & Sejnowski, 1995). Data were then re-referenced to the average of the two mastoid sensors. Data segments corresponding to the timing and duration of the two stories were extracted and downsampled to 100 Hz.

#### 2.4.2 Predictor variables

In order to measure the neural representations of speech at different levels of processing, multiple predictor variables were generated. Each predictor variable is a continuous time-series, aligned with the stimulus, which quantifies a specific feature, hypothesized to evoke a neural response (see Figure 1 for an overview). The predictors for auditory and English linguistic processing closely followed previously used representations that were developed as measures of processing English as a native language (see Brodbeck et al., 2022).

**Auditory processing** was assessed using an auditory spectrogram and acoustic onsets. **Linguistic processing** was assessed at the sublexical, word-form, and sentence level using information-theoretic models. These models are all predictive language models that predict upcoming speech phoneme-by-phoneme, but they differ by taking into account different amounts of context (for a detailed theoretical motivation see Brodbeck et al., 2022). Previous research has shown that such models track speech comprehension more closely than acoustic models (Brodbeck et al., 2018; Verschueren et al., 2022). **Sublexical processing** was assessed using a context that consisted of a sublexical phoneme sequence, taking into account only the previous 4 phonemes. **Word form processing** was assessed using a within-word context, taking into account only the phonemes in the current word. **Sentence level processing** was assessed using a multi-word context consisting of the preceding four words. At all linguistic levels, the influence of context representations on brain responses was operationalized through *phoneme surprisal* Equation 1 and *phoneme entropy* Equation 2 measures:

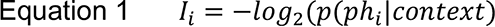

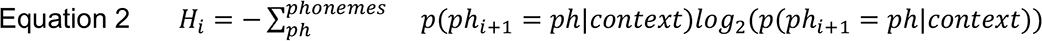

*Phoneme surprisal* at position *i*, *I*_*i*_, reflects how surprising the phoneme at position *i* is, given a certain context (e.g., sublexical phoneme surprisal quantifies how surprising the current phoneme is based on a prediction using the past 4 phonemes; sentence level phoneme surprisal reflects how surprising the current phoneme is based on a prediction using the past four words and the current partial word). *Phoneme entropy H*_*i*_ reflects how much uncertainty there is about the identity of the next phoneme. For lexical processing models, *cohort entropy* Equation 3 additionally reflects how much uncertainty there is about what the current word is:

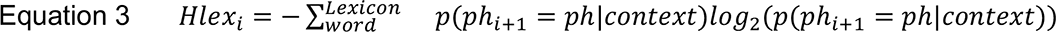

This again, depends on what context is used. For example, using only the word-form context, *s…* is much more uncertain than when using the sentence context, *coffee with milk and s…*. Significant brain responses related to these variables were taken as indicators of incremental linguistic processing of speech at these different levels. Finally, in addition to information-theoretic models, neural correlates of lexical segmentation were controlled for using a predictor for responses to word onsets (Brodbeck et al., 2018). A predictor with an equally scaled impulse at each phoneme onset was included to control for any phoneme-evoked response not modulated by the predictors of interest (analogous to the intercept term in a regression model).

To generate the sublexical and lexical predictors, word- and phoneme locations are needed, which were determined in the auditory stimuli using forced alignment. To that end, an English pronunciation dictionary was defined based on merging the Montreal Forced Aligner (McAuliffe et al., 2017) English dictionary with the Carnegie-Mellon Pronouncing Dictionary, and manually adding five additional words that occurred in the short story. The time point of words and phonemes in the acoustic stimuli were then determined using the Montreal Forced Aligner. Below, the different predictors and how they were created are explained in detail.

##### 2.4.2.1 Auditory processing

Two predictors were used to assess (and control for; Daube et al., 2019; Gillis et al., 2021) auditory representations of speech: An auditory spectrogram and an acoustic onset spectrogram. The auditory spectrogram reflects moment by moment acoustic power, using a transformation approximating peripheral auditory processing, and thus models sustained neural responses to the presence of sound. The onset spectrogram specifically contains acoustic onset edges, and thus models transient response to the onset of acoustic features.

An auditory spectrogram with 128 bands ranging from 120 to 8000 Hz in ERB space was computed at 1000 Hz resolution with the gammatone library (Heeris, 2018). The spectrogram was log-transformed to more closely reflect the auditory system’s dynamic range. For use as the auditory spectrogram predictor variable, the number of bands was reduced to 8 by summing 16 consecutive bands.

The 128 band log spectrogram was transformed using a neurally inspired auditory edge detection algorithm (Fishbach et al., 2001) to generate the acoustic onset spectrogram (Brodbeck et al., 2020). For use as a predictor variable, the number of bands was also reduced to 8 by summing 16 consecutive bands.

##### 2.4.2.2 Sublexical English representations

Sublexical representations were assessed using a context consisting of phoneme sequences. To that end, first a probabilistic model of phoneme sequences in English without consideration of word boundaries was generated: all sentences of the SUBTLEX-US corpus (Brysbaert & New, 2009) were transcribed to phoneme sequences by substituting each word with its pronunciation from the pronunciation dictionary and concatenating these pronunciations across word boundaries. The resulting phoneme strings were used to train a 5-gram model using KenLM (Heafield, 2011). This 5-gram model was then used to estimate probability distributions for the next phoneme at each position in the story (*p*(*ph*_*i*\*+_|*ph*_*i-3*_, *ph*_*i-2*_, *ph*_*i-1*_, *ph*_*i*_), with *i* indexing the current position in the story). These probability distributions were used to generate two predictors, *phoneme surprisal I*_*i*_ and *phoneme entropy H*_*i*_ (Equations Equation 1 and Equation 2). Each of these predictors was constructed by placing an impulse at the onset of each phoneme, scaled by the respective *surprisal* or *entropy* value. These predictors were used to measure the use of sublexical phonotactic knowledge during speech processing.

Additionally, a *phoneme onset* predictor was included, with impulse size of one at each phoneme, to serve as an intercept for the sublexical predictors (i.e., capturing any response that occurs to phonemes but is not modulated by any of the quantities of interest).

##### 2.4.2.3 English word-form representations

A *word onset* predictor was generated with equal sized impulses at each word onset to assess lexical segmentation (Brodbeck et al., 2018; Sanders et al., 2002). This predictor was taken as an indicator of lexical segmentation, when contrasted with the phoneme predictor which measures responses related to phonetic processing without regard for lexical segmentation.

Word-form representations were assessed using a model of word recognition that takes into account word boundaries, but disregards the preceding multi-word context. This model is based on the cohort model of word recognition (Marslen-Wilson, 1987). A lexicon was defined based on the pronunciation dictionary (also used for forced alignment), in which each unique grapheme sequence identifies a word, and each word may have multiple pronunciations. At each word boundary, the cohort is initialized using the whole lexicon, with the prior likelihood for each word proportional to its frequency in the SUBTLEX-US corpus (Brysbaert & New, 2009). At each phoneme position, the cohort is pruned by removing all words whose pronunciations are incompatible with the new phoneme, and word likelihoods are renormalized. Thus, at the *j*th phoneme of the *k*th word, this cohort model tracks the probability distribution over what word the current phoneme sequence could convey as *p*(*word*_*k*_|*ph*_1_, . . ., *ph*_*j*_). Since each word is associated with a likelihood and also makes a prediction about what the next phoneme would be, this amounts to a predictive model for the next phoneme *p*(*ph*_*j* + 1_|*ph*_1_, . . ., *ph*_*j*_). These evolving probability distributions over the lexicon are in turn used to compute *phoneme surprisal* (i.e., how surprising the current phoneme is given what words were still in the cohort at the previous position), *phoneme entropy* (uncertainty about the next phoneme) and *lexical entropy* (uncertainty about what the current word is) (Equations Equation 1, Equation 2 and Equation 3). These predictors were used to measure word-form processing independent of the wider sentence context. Thus, if sublexical surprisal is a significant predictor of brain activity, this suggests that listeners use sublexical phoneme sequences to make predictions about upcoming phonemes; if word-form surprisal is significant, this suggests that they also use information about what the current word could be.

##### 2.4.2.4 Sentence level representations

Sentence-level processing was assessed using a lexical model augmented by the preceding multi-word context. The model is identical to the English word-form model, except that now in the word-initial cohorts, prior probabilities for the words are not initialized based on their lexical frequency, but instead based on a case-insensitive, lexical 5-gram model (Heafield, 2011) trained on the word sequences in the SUBTLEX-US corpus (Brysbaert & New, 2009). Thus, instead of tracking the probability of a word k, given the phonemes of word k heard so far, *p*(*word*_*k*_|*ph*_1_, . . ., *ph*_*j*_), this model tracks the probability of a word k given the previous 4 words in addition to the phonemes of word k, *p*(*word*_*k*_|*word*_*k* - 4_, . . ., *word*_*k* - 1_, *ph*_1_, . . ., *ph*_*j*_). Predictors based on this language model were used to measure the use of the multi-word context during speech processing.

##### 2.4.2.5 Sublexical Dutch representations

Interference from Dutch sublexical phonotactic knowledge was assessed analogous to the English sublexical model, but trained on Dutch lexical statistics. Phoneme sequences were extracted from version 2 of the Corpus Gesproken Nederlands (CGN; Oostdijk et al., 2002), and used to train a phoneme 5-gram model (Heafield, 2011). Since Dutch and English have different phoneme inventories, and the 5-gram model was trained on Dutch phonemes, each English phoneme of the stimulus story was transcribed to the closest Dutch phoneme. The resulting phoneme sequence, reflecting a transcription of the English story with the Dutch phoneme inventory, was then used to compute *phoneme surprisal* and *phoneme entropy* as for the sublexical English model using the phoneme 5-gram model trained on Dutch. The resulting predictors were used to measure brain responses that would indicate that listeners activated their knowledge of their native Dutch sublexical phonotactics when listening to the English story.

##### 2.4.2.6 Word-level native language interference

To test for interference from native language word knowledge, we generated two alternative word-form models. These were built and used like the English word-form model, differing only in the set of lexical items that were included in the pronunciation lexicon. First, we built a Dutch word-form model (*word-form*D). This model contained only Dutch words and their pronunciations, taken from the CGN lexicon. In order to evaluate lexical cohorts in the (English) input phoneme inventory, Dutch phonemes were mapped to the closest available English equivalent (as for the sublexical Dutch model), or, in the absence of a close English phoneme, to a special out-of- inventory token (which always leads to exclusion from the cohort when encountered). Relative lexical frequencies were taken from the SUBTLEX-NL corpus (Keuleers et al., 2010) to closely match the way in which the English lexical frequencies were determined using SUBTLEX-US. Finally, we also built an English/Dutch combined lexicon, using the union of the two pronunciation dictionaries (*word-form*ED).

#### 2.4.3 mTRF analysis

An mTRF is a linear mapping from a set of *nx* predictor time series, *xi,t*, to a response time series *yt*. The response at time *t* is predicted by convolving the predictors with a kernel *h*, called the mTRF, at a range of delay values *τ*:

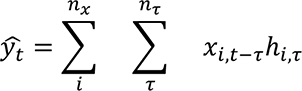

The mTRFs were estimated using the boosting algorithm (David et al., 2007) implemented in Eelbrain (Brodbeck et al., 2021), separately for each story. Delay values (*τ*) ranged from -100 to 850 ms. For 5-fold cross-validation, predictors and EEG responses were split into 5 segments of equal length. To predict the EEG response to each segment, an mTRF was trained on the four remaining segments. This mTRF in turn was the average of 4 mTRFs, which were trained by iteratively using one of the four segments as validation data and the remaining 3 segments as training data. The mTRFs were trained using coordinate descent to minimize ℓ2 error of the predicted response in the training data. If after any training step the ℓ2 error in the validation data increased, then this last step was undone and the TRF corresponding to this predictor was frozen (i.e., excluded from further modification by the fitting algorithm). Fitting continued until all TRFs were frozen.

##### 2.4.3.1 Predictive power

Evidence for specific neural representations was assessed by testing whether the corresponding predictors significantly contributed to predicting the held-out EEG data. In order to evaluate the predictive power of a specific predictor, or a group of predictors, two mTRFs were estimated: one for the full model (i.e., all predictors), and one for the full model minus the predictor(s) under investigation. Since the predictive power was measured on data that was not seen during model estimation, the null hypothesis is that the two models predict the data equally well, whereas the alternative hypothesis is that adding the predictor(s) under investigation improves the model fit.

Predictive power was quantified as the proportion of the variance explained in the EEG data. This was calculated as 1 − ∑_*t*_ (*y*_*t*_ − *y*_*t*_)^2^/∑_*t*_ *y*_*t*_^2^. In order to test whether the predictive power of two models differed reliably across participants, we first compared the predictive power of the two models, averaged across all sensors, with a repeated measures *t*-test. We report Cohen’s *d* effect sizes. In case there was a significant difference, we then used mass-univariate tests to find sensor regions that contributed to the effect. These mass-univariate tests were cluster-based permutation tests (Maris & Oostenveld, 2007), using as cluster-forming threshold a *t*-value corresponding to an uncorrected *p*=.05, and estimating corrected *p*-values for each cluster’s *cluster-mass* statistic (summed *t*-values) on a null distribution estimated from 10,000 random permutations of condition labels.

In some comparisons where we are interested in the null hypothesis (e.g., whether there is evidence for native language interference in a given condition) we also report Bayes factors (*B*) (Rouder et al., 2009) estimated using the BayesFactor R library, version 0.9.12-4.4 (Morey et al., 2022). For directional contrasts (e.g., that predictive power is > 0), we report the Bayes factor for evidence in favor of the value being >0 vs <0 (Morey & Rouder, 2011).

##### 2.4.3.2 Correlations with language proficiency

To test whether language proficiency measures explained neural responses, we analyzed the predictive power of the different language models as a function of the LexTale and LLAMA_D test scores. As dependent measure we extracted the predictive power for a given set of predictors across all EEG sensors. This measure of predictive power is the difference in explained variance

(*Δv*) between two models which differ only in the inclusion or exclusion of the predictors under investigation. We then analyzed the predictive power in R (R Core Team, 2021) using linear mixed effects models as implemented in lme4 (Bates et al., 2015), with the following formula:

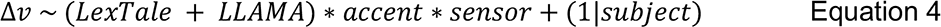

Including higher level random effect structure generally resulted in singular fits, with one exception: for the analysis of auditory responses, we were able to specify *sensor* as random effect. We tested for significant effects using likelihood ratio tests. In order to minimize the number of comparisons, we first tested whether there was any effect of proficiency, by comparing model Equation 4 with a model in which all terms including LexTale were removed (and analogous for aptitude/LLAMA). In case of a significant difference, we then tested whether the effect of proficiency was modulated by speaker accent by comparing model Equation 4 to a model lacking only terms including a LexTale:accent interaction. When significant interactions with *accent* were detected we fit separate linear models for the English and Dutch accented conditions.

When we detected significant effects involving *LexTale* or *LLAMA*, we then performed further analyses to explore the topographic distribution of these effects across EEG sensors. For this, we fitted a multiple linear regression with the following model, independently at each sensor and for each accent condition:

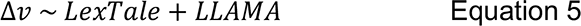

We show topographic plots of the *t* statistic corresponding to the predictors of interest from this regression (LexTale/LLAMA). We further selected sensors at which *t*≥2 to produce scatter plots for illustrating the relationship, and for analyzing TRF magnitudes (next paragraph).

We further analyzed the TRFs corresponding to the predictors that were related to proficiency, to gain more insights in the brain dynamics underlying the predictive power effects. If a predictor contributes to the predictive power of a model, it does so through the weights in its TRF. We investigated these weights to gain more insight into the time-course at which the predictor’s features affect the brain response. For this, we upsampled TRfs to 1000 Hz and calculated the TRF magnitude as a function of time (for each lag, the sum of absolute values of the weights across sensors and, for acoustic predictors, frequency). We analyzed these time-courses using a mass-univariate multiple regression model with the same model as in Equation 5, correcting for multiple comparisons across the time course (0–800 ms) with cluster-based permutation tests with the same methods described for the analysis of predictive power.

## 3 Results

We hypothesized that acquiring a new language involves learning new acoustic-phonetic representations, as well as developing predictive language models that use different contexts to anticipate upcoming speech. Here we looked for evidence of such representations in EEG responses to narrative speech. To address the research questions outlined in the Introduction, we proceeded in three steps: 1) we verified that the previously described predictive language models for English at the sublexical, word-form and sentence level (Brodbeck et al., 2022; Z. Xie et al., 2023) were also significant predictors for EEG responses of non-native listeners; 2) we tested the influence of Dutch, the native language, on processing of English by testing the predictive power of language models that incorporate Dutch language statistics; 3) we determined to what extent these effects are modulated by English proficiency (LexTale) and acoustic-phonetic aptitude (LLAMA_D).

### 3.1 Proficiency and aptitude test results

Figure 2 shows that English proficiency (LexTale) and acoustic-phonetic aptitude (LLAMA_D) were uncorrelated (*r*(37)=-.06, *p*=.700). This confirms that the two tests measure independent aspects of second language ability.

**Figure 2.**
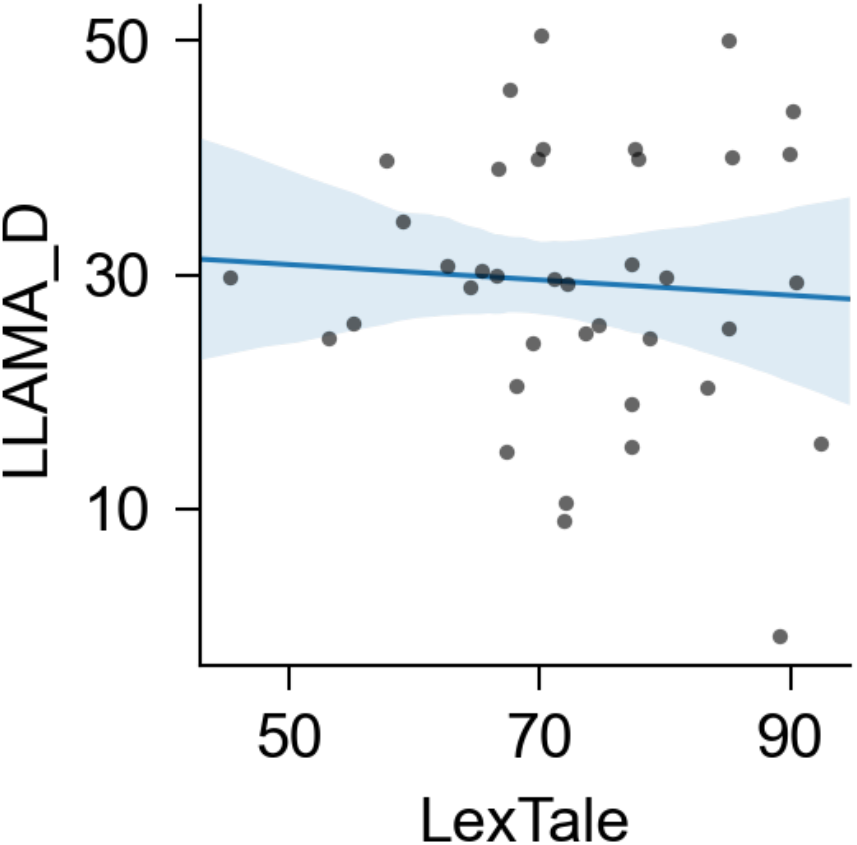
LexTale and LLAMA_D measure independent aspects of language ability. Each dot represents scores from one participant. The line represents the linear fit, with a 95% confidence interval estimated from bootstrapping. Because scores take discrete values, a slight jitter was applied to the data for visualization after fitting the regression.

### 3.2 Robust acoustic and linguistic representations of the non-native language

To test whether listeners formed a specific kind of representation, we tested whether a predictor designed to capture this representation has unique predictive power, i.e., whether an mTRF model including this predictor is able to predict held-out EEG responses better than the same model but without the specific predictor. We initially started with a model containing predictors for auditory and linguistic representations established by research on native language processing (Brodbeck et al., 2022), illustrated in Figure 1:

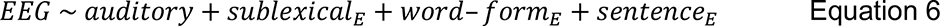

The *auditory* predictors consisted of an auditory spectrogram and onset spectrogram. The English (E) linguistic predictors were based on three information-theoretic language models, all modeling incremental, phoneme-by-phoneme information processing: a *sublexical* phoneme sequence model, a *word-form* model and a *sentence* model.

To determine whether the different components of model Equation 6 describe independent neural representations, we tested for each component whether it significantly contributed to the predictive power of the full model (Figure 3 and Table 1). We first tested the average predictive power in the two stories (American & Dutch, A&D, Figure 3 first row), then tested for a difference between the two stories (American vs Dutch, AvD, not shown in Figure 3) and confirmed the effect separately in the American (A) and Dutch (D) accented stories (Figure 3 second and third row). Auditory predictors (Figure 3-A) and the three language levels (Figure 3-B) all made independent contributions to the overall predictive power, and none of them differed between stories (statistics in Table 1). The topographies of predictive power are comparable to known distributions reflecting auditory responses, suggesting contributions from bilateral auditory cortex (e.g. Lütkenhöner & Mosher, 2007), similar to native listeners’ responses (Brodbeck et al., 2022). Taken together, these results suggest that non-native Dutch listeners, as a group, use English sublexical transition probabilities (sublexical context), word-form statistics (word-form context), as well as multi-word transition probabilities (sentence context) to build incremental linguistic representations when listening to an English story.

**Figure 3.**
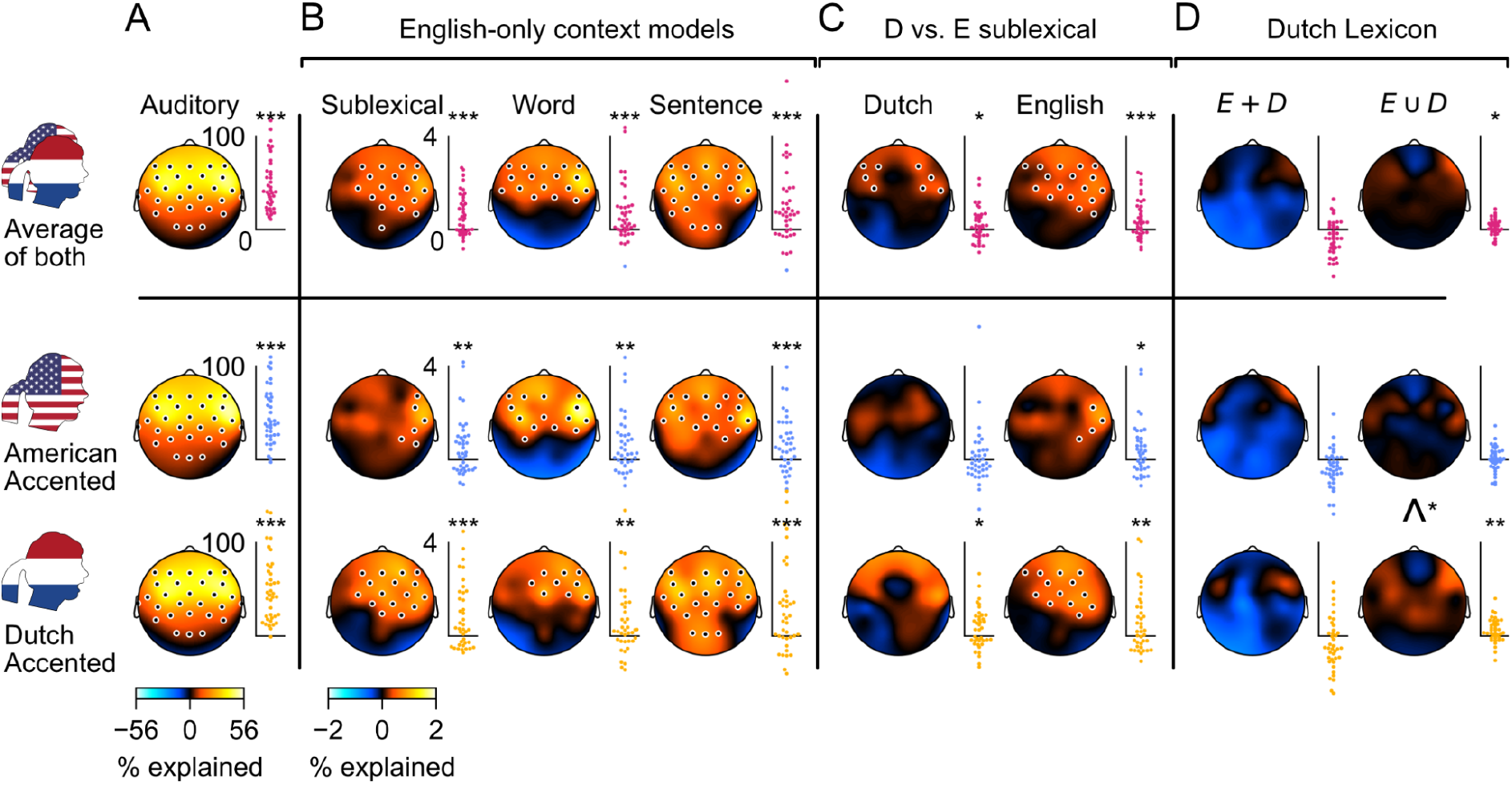
Auditory and linguistic neural representations in Dutch listeners when listening to an English story. Each swarm-plot shows the change in predictive power for held-out EEG responses when removing a specific set of predictors (each dot represents the change in predictive power, averaged across sensors, for one participant). Predictive power is expressed in percent of the variance explained by the English model (model Equation 6) averaged across subjects. Stars indicate significance based on a one-tailed related measures *t*-test. Topographic maps show corresponding sensor-specific data, with predictive power expressed as percent of model Equation 6 at the best sensor. The marked sensors form significant clusters in a cluster-based permutation test based on one-tailed *t*-tests. (A) Auditory predictors contribute a large proportion of the explained variance. The measure is based on the difference in predictive power between the English model Equation 6, and a model missing auditory predictors (acoustic onset and auditory spectrogram). (B) All three linguistic models significantly contributed to the predictive power of the English model, in both stories. Note that predictive power can be negative, indicating that adding the given predictor made cross-validated predictions worse. (C) A sublexical Dutch model, reflecting phoneme sequence statistics in Dutch (*sublexicalD*), significantly improved predictions even after controlling for English phoneme sequence statistics (*sublexicalE*), suggesting that Dutch listeners create expectations for phoneme sequences that would be appropriate in Dutch even when listening to English. The English sublexical model remained significant after adding the Dutch sublexical model. (D) Addition of Dutch word-forms suggests word recognition with competition from a combined lexicon: Adding a word-form model using only Dutch pronunciations (*word-form*D) does not improve predictions (left column, comparison: model Equation 8 > Equation 7), suggesting that native language word recognition does not proceed in parallel. In contrast, replacing the English word-form model *word-form*E with a merged word-form model *word-form*ED, which combines English and Dutch word-forms, leads to improved predictions of EEG responses to Dutch accented speech (right column, comparison: Equation 9 > Equation 7). *: *p*≤.05; **: *p*≤.01; ***: *p*≤.001.

**Table 1.** Statistics for the predictive power of English language models, averaged across all EEG sensors (corresponding to swarm plots in Figure 3). Significant results in bold font (p≤.05).

|  | American & Dutch |  |  | American vs Dutch |  |  | American |  |  | Dutch |  |  |
| --- | --- | --- | --- | --- | --- | --- | --- | --- | --- | --- | --- | --- |
|  | *t*(38) | *p* | *d* | *t*(38) | *p* | *d* | *t*(38) | *p* | *d* | *t*(38) | *p* | *d* |
| Auditory | **10.7** | **< .001** | **1.71** | -0.37 | .716 | -0.06 | **9.87** | **< .001** | **1.58** | **8.86** | **< .001** | **1.42** |
| Sublexical | **4.68** | **< .001** | **0.75** | -0.87 | .387 | -0.14 | **2.81** | **0.004** | **0.45** | **3.65** | **< .001** | **0.59** |
| word-form | **3.95** | **< .001** | **0.63** | 0.7 | .485 | 0.11 | **3.11** | **0.002** | **0.50** | **2.69** | **.005** | **0.43** |
| Sentence | **4.31** | **< .001** | **0.69** | -0.83 | .411 | -0.13 | **3.39** | **< .001** | **0.54** | **3.75** | **< .001** | **0.60** |

Previous results suggested that native English listeners activate sublexical, word-form and sentence models in parallel, evidenced by simultaneous early peaks in their brain response to phoneme surprisal (Brodbeck et al., 2022). This contrasts with an alternative hypothesis of cascaded activation, which would predict that lower level models are activated before higher level models, i.e., first the sublexical, then the word-form, and then the sentence model (e.g. Zwitserlood, 1989). Figure 4 shows TRFs for phoneme surprisal associated with the three language models in model Equation 6. Each language model is associated with an early peak around 60 ms latency (peaks might appear earlier than expected because the forced aligner does not account for coarticulation). This suggests that the different language models are activated in parallel in non-native listeners, as they are in native listeners.

**Figure 4.**
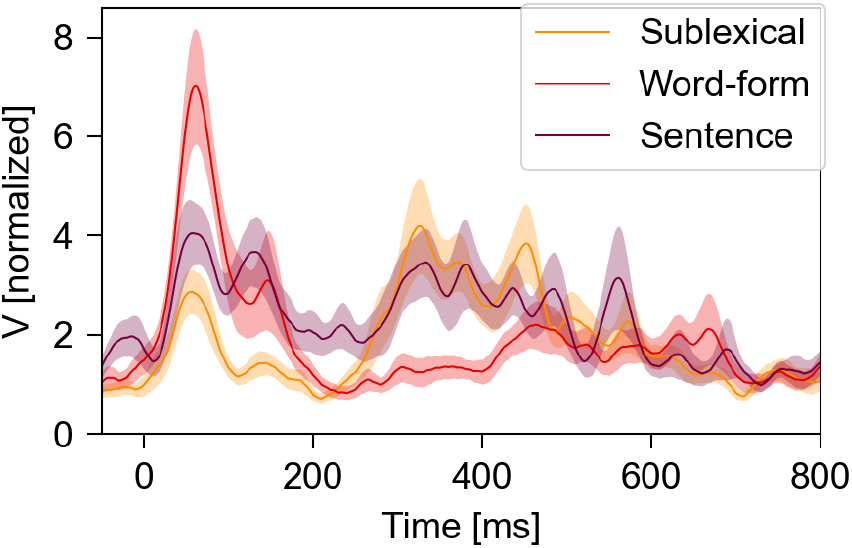
Simultaneous early peaks in temporal response functions (TRFs) suggest parallel processing. Each line represents the TRF magnitude (sum of the absolute values across sensors). The y-axis represent original values ✕10^-3^.

**Figure 5.**
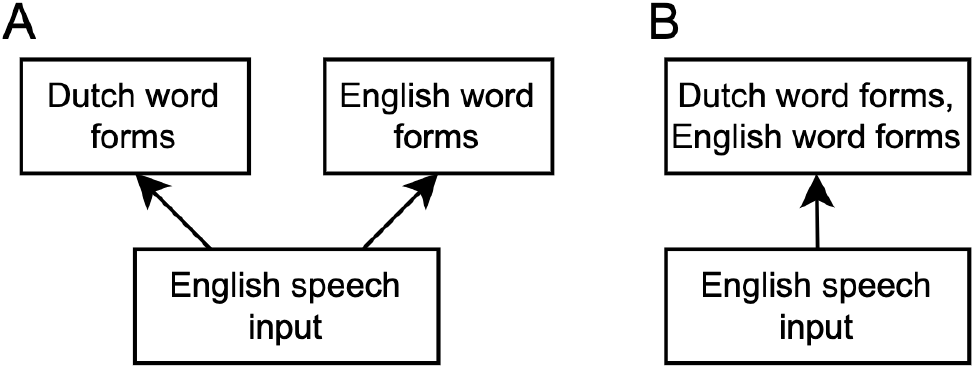
Alternative explanations for activation of native language lexical candidates during non-native listening. (A) The native language and the non-native language lexicons are independent systems that are both activated in parallel by acoustic input. Outputs of the two systems may still interact, e.g. in guiding eye movements in visual world studies. (B) Word-forms from both languages compete in a single recognition system.

### 3.3 Influence of the native language on non-native language processing

#### 3.3.1 Dutch sublexical phoneme sequence model is activated when listening to Dutch-accented English

Learning English as a non-native language entails acquiring knowledge of the statistics of English phoneme sequences, i.e., a new sublexical context model. Given the relatively large overlap of the Dutch and English phonetic inventories, the native language Dutch sublexical model might still be activated when listening to English. To test whether this is indeed the case, we added predictors from a Dutch sublexical context model, *sublexicalD*, to model Equation 6. The Dutch sublexical model was analogous to the English sublexical model, containing phoneme surprisal and entropy based on Dutch phoneme sequence statistics. To test whether the Dutch sublexical model can explain EEG response components not accounted for by the English sublexical model, the predictive power of the model containing both, model Equation 7, was compared to a model without the Dutch sublexical predictor Equation 6, and vice versa.

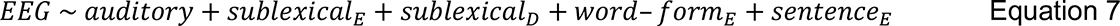

When averaging the predictive power of the two stories, both the Dutch and the American sublexical models contributed explanatory power (Figure 3-C; *sublexical*D: *t*(38)=1.72, *p*=.046, *d*=0.28; *sublexical*E: *t*(38)=3.67, *p*<.001, *d*=0.59). The explanatory power of the English *sublexical*E model remained robust across stories (A: *t*(38)=2.30, *p*=.013, *d*=0.37; D: *t*(38)=2.87, *p*=.003, *d*=0.46; AvD: *t*(38)=-0.67, *p*=.506, *d*=-0.11). However, this was not the case for the Dutch *sublexical*D model. Evidence for an effect of the Dutch *sublexical*D model in the Dutch-accented story was strong (*t*(38)=2.32, *p*=.013, *d*=0.37, *B*=64.45). However, evidence for interference in the American-accented story was weak, with only negligible evidence in favor of some interference (*t*(38)=0.33, *p*=.373, *d*=0.05, *B*=1.66). The effect was not significantly stronger in the Dutch accentend story compared to the American accented story, suggesting that some caution is warranted, but the Bayes factor suggests some evidence in favor of a stronger effect in the Dutch accented story (*t*(38)=1.02, *p*=.156, one-tailed, *d*=0.16, *B*=5.12). We conclude that interference was likely stronger in the Dutch accented story, but some interference may have occurred in both stories.

#### 3.3.2 Dutch and English word forms are activated together when listening to Dutch accented English

Several previous studies suggest that Dutch word forms are activated alongside English word forms when listening to English (see Introduction). This could occur in two different ways (Figure 5): Dutch word forms could be activated in a separate lexical system, without competing with English word forms. Alternatively, Dutch and English word forms could compete for recognition in a connected lexicon.

To test the first possibility (Figure 5-A), we tested whether a separate word-form model with only Dutch word forms, *word-formD*, improved predictive power when added in addition to the English word-form model:

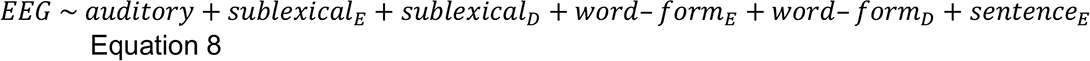

This implements the hypothesis that two independent brain systems track English and Dutch word forms independently, i.e., at each phoneme the two systems encounter different amounts of surprisal and entropy according to their respective lexicon, and each system generates a neural response, with the two responses combining in an additive manner. Comparing model Equation 8 with Equation 7 tests for the existence of such a Dutch lexical model alongside the English model. The results showed that the Dutch word-form model did not further improve predictions after controlling for other predictors. Indeed, the addition of the Dutch word-form model made predictions worse, as might be expected in cross-validation from a predictor that adds noise (A&D: *t*(38)=-3.09, *p*=.998; AvD: *t*(38)=-0.20, *p*=.840).

To test the second possibility (Figure 5-B), we tested a merged lexicon, i.e., a model analogous to the English word-form model, but including both English and Dutch word forms: *word-formED*. This merged word-form model embodies the hypothesis that a single lexical system detects word forms of both languages, i.e., at each phoneme there is only a single surprisal and entropy value, which depends on the expectation that the current word could be English as well as Dutch. Since this merged word-form model is hypothesized as an alternative to the English-only word-form model (*word-formE*), we here tested the effect on predictive power of substituting the merged word-form model for the English word-form model (two-tailed test) – i.e., we compared model Equation 9 with Equation 7:

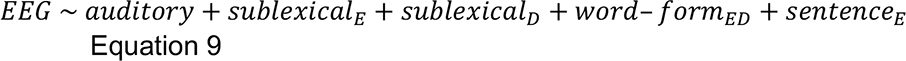

Overall, the merged word-form model improves predictions over the English word-form model (A&D: *t*(38)=2.39, *p*=.022, two-tailed, *d*=0.38; Figure 3-D). It is conceivable that compared to the American accent, a Dutch accent, which better matches Dutch phonological categories, increases activation of Dutch competitors. Indeed, when analyzing accents separately, the evidence in favor of the merged *word-form*ED model was strong in the Dutch-accented story (*t*(38)=2.98, *p*=.005, *d*=0.48, *B*=312.26) and negligible in the American accented story (*t*(38)=0.29, *p*=.771, d=0.05, *B*=1.57), and there was considerable evidence for a difference between speaker accents (AvD: *t*(38)=1,77, *p*=.042, one-tailed, *d*=0.28, *B*=20.11).

Crucially, the merged word-form model was significantly better than the parallel lexicon model, confirming that a lexicon with direct lexical competition between candidates from the two languages better accounts for the data than activation in two parallel lexica (model Equation 9 vs. Equation 8, A&D: *t*(38)=4.11, *p<*.001; A: *t*(38)=2.98, *p*=.005; D: *t*(38)=3.10, *p*=.004).

### 3.4 Modulation of non-native language processing by language proficiency

We next asked whether the acoustic and linguistic representations are modulated by non-native language proficiency. We used model Equation 9 as the basis for these analyses, because the results reported above suggested that Equation 9 was the best model. Thus, predictive power reported in the following section was always calculated by removing the relevant predictors from model Equation 9. We used linear mixed effects models to determine whether a given representation is influenced by language proficiency (LexTale) or language learning aptitude (LLAMA_D), and if so, whether this relationship is modulated by speaker accent. Table 2 shows results for the LexTale score, and Table 3 shows corresponding results for the LLAMA_D score.

**Table 2.** Influence of proficiency on the predictive power of different EEG model components, determined with linear mixed effects models. The *LexTale* column reports tests for any influence of lexTale, i.e., whether all terms including LexTale combined (main effect and interactions) significantly improved models (likelihood ratio tests). If this was the case, the *LexTale × accent* column reports whether the effect of LexTale was modulated by speaker accent by testing whether the terms including a LexTale:accent interaction significantly improved models. Reported *p*-values are uncorrected; results in bold indicate significance (*p*<.05) after correcting for false discovery rate among the tests reported in this table.

|  | LexTale |  | LexTale × accent |  |
| --- | --- | --- | --- | --- |
| | $\chi^2(28)$ | *p* | $\chi^2(28)$ | *p* |
| Auditory | **123.41** | **<.001** | **48.70** | **.009** |
| Sublexical<sub>E</sub> | **156.87** | **<.001** | **135.82** | **<.001** |
| Sublexical<sub>D</sub> | 59.00 | .367 |  |  |
| Word-form<sub>ED</sub> | 58.78 | .374 |  |  |
| Word-form<sub>ED>E</sub> | 63.53 | .228 |  |  |
| Sentence | 46.99 | .799 |  |  |

**Table 3.**
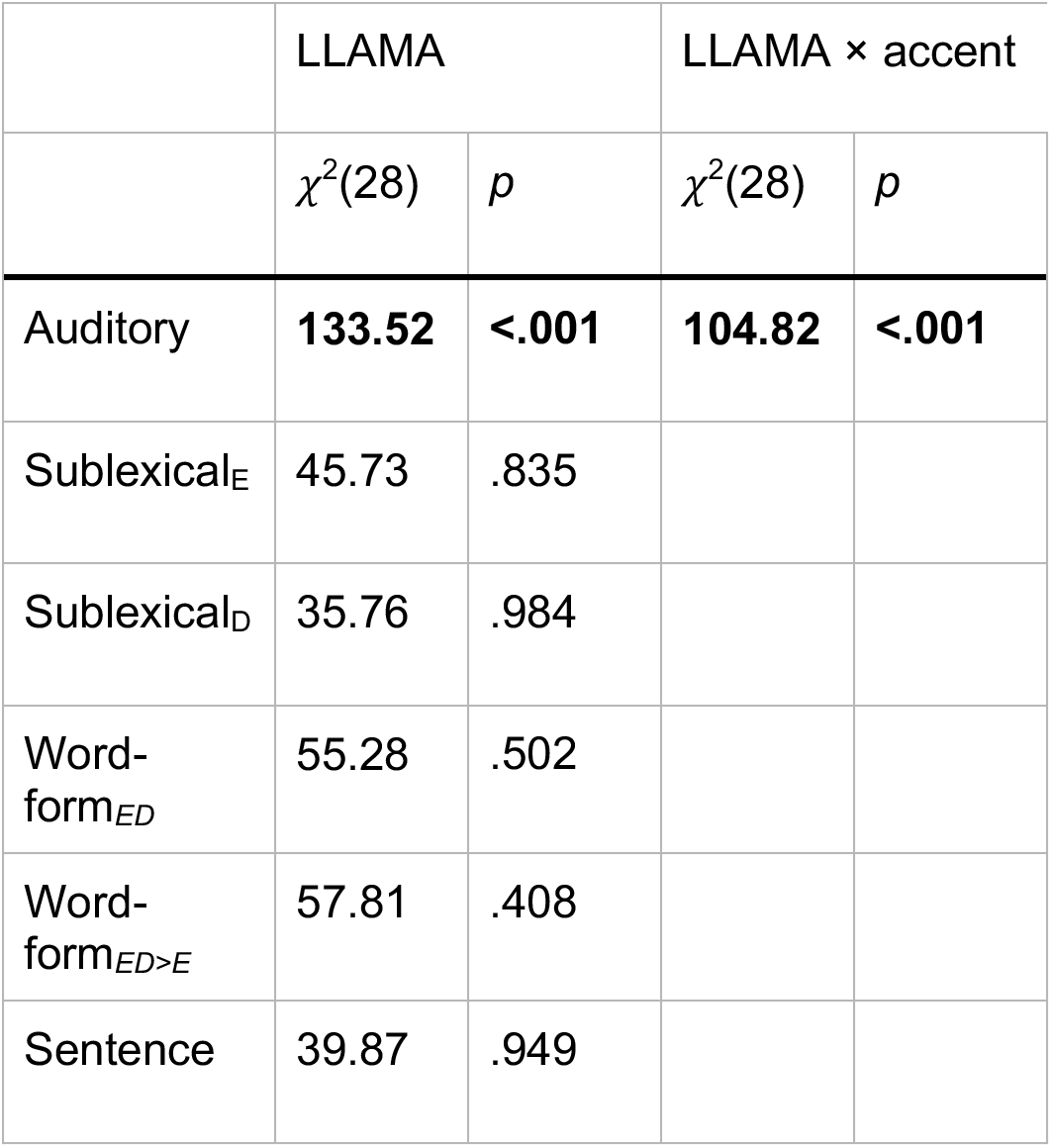
Influence of acoustic-phonetic aptitude as measured by the LLAMA_D test on the predictive power of different EEG model components. Details as in Table 2.

#### 3.4.1 Increased proficiency (LexTale) is associated with reduced late auditory responses

The predictive power of the auditory predictors was significantly modulated by proficiency as measured by LexTale (Table 2). Even though this association differed between accents, it was independently significant for American and Dutch accented speech (A: *χ*^2^(28)=59.59, *p*<.001; D: *χ*^2^(28)=98.25, *p*<.001). In both cases, individuals with higher proficiency had weaker auditory representations, and this modulation involved electrodes across the head (Figure 6-AC). An analysis of the TRFs suggests that in both accent conditions, lower proficiency was associated with larger sustained auditory responses at relatively late lags (A: 250–393 ms, *p*<.001; D: 220– 287 ms, *p*=.008 and 348–407 ms, *p*=.015; Figure 6-BD, right panels). These results indicate that listeners with lower proficiency exhibit enhanced sustained auditory representations at relatively late lags.

**Figure 6.**
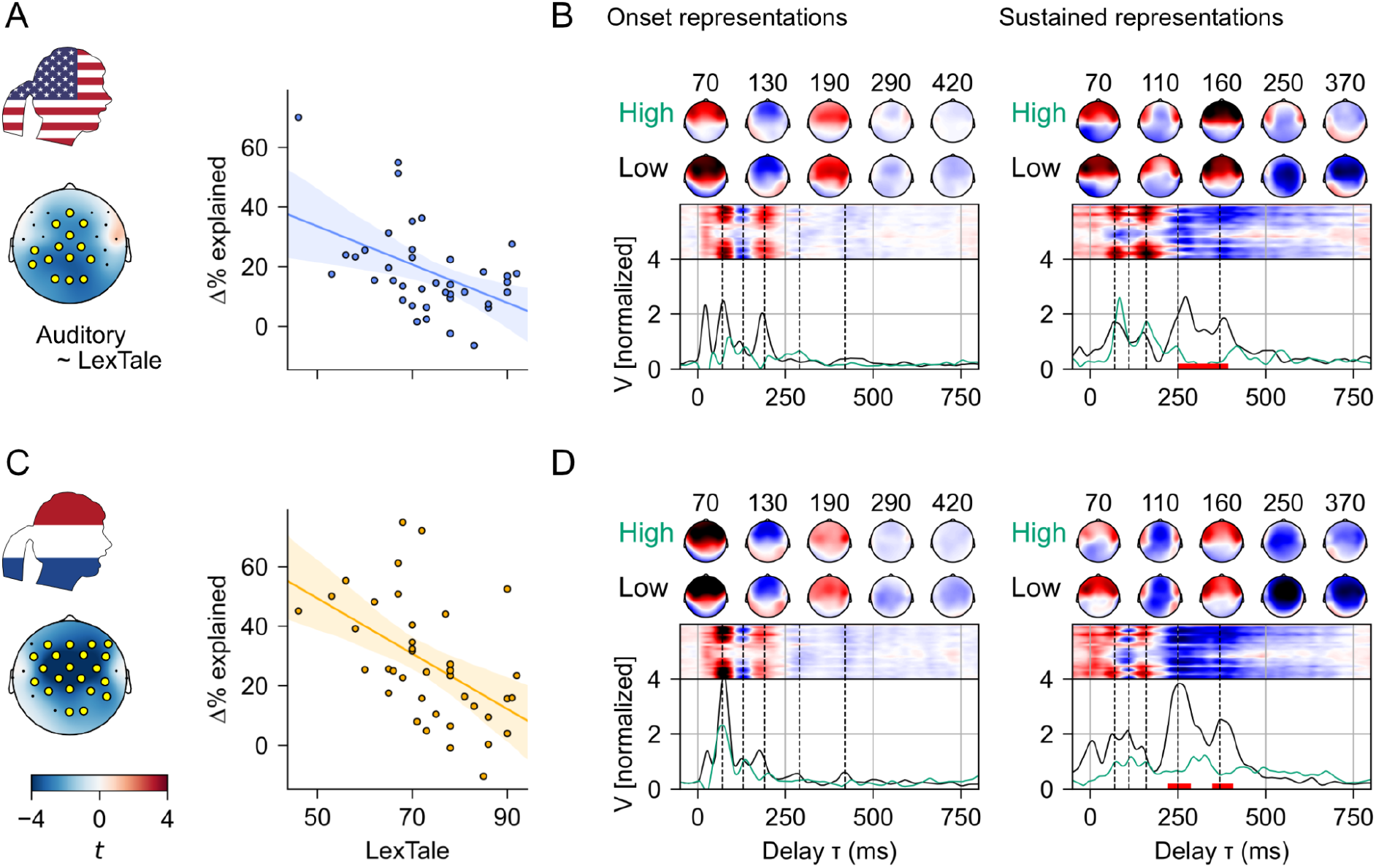
Auditory responses are modulated by non-native language proficiency. (A) The strength of the auditory responses to American-accented English decreases with increased language proficiency. The topographic map shows the multiple linear regression *t* statistic, for the influence of LexTale scores on the predictive power of the auditory model, when the speaker had an American accent. Sensors with *t* values exceeding 2 (positive or negative) are marked with yellow. The scatter-plot shows the predictive power of the auditory model (y-axis, average of marked sensors) against LexTale scores (x-axis). Each dot represents one participant. The solid line is a direct regression of predictive power on the LexTale score; bands mark the 95% confidence interval determined by bootstrapping. (B) TRFs suggest that less proficient listeners have stronger sustained auditory representations at later response latencies. The line-plot shows the magnitude of the TRF across sensors as predicted by the multiple regression for small and large values of LexTale (60, 90), while keeping other regressors at their mean. Red bars at the bottom indicate a significant effect of LexTale (regression model Equation 5). The rectangular image plot above shows the average TRF for each sensor, and the topographic maps show specific time points (marked by dashed black lines below) for participants with low and high LexTale scores (median split). While auditory TRFs were estimated as mTRFs for 8 spectral bands in each representation, for easier visualization and analysis the band-specific TRFs were summed across bands (after calculating magnitudes where applicable). (C) The strength of auditory responses to Dutch- accented English also decreases with increased language proficiency. (D) TRFs to Dutch- accented speech show a similar effect of proficiency on sustained representations as in American accented speech.

#### 3.4.2 Acoustic-phonetic aptitude (LLAMA_D) is associated only with processing of Dutch-accented speech

The predictive power of auditory responses was also modulated by acoustic-phonetic aptitude, and this effect was qualified by an interaction with speaker accent (Table 3). Figure 7-AC illustrate the pattern creating this interaction. Acoustic responses to the American accented story were not modulated by aptitude (*χ*^2^(28)=17.74, *p*=.932), but responses to the Dutch accented story were (*χ*^2^(28)=88.75, *p*<.001), with a broadly distributed topography (Figure 7-C). Consistent with this, TRF magnitudes were not related to phonetic ability in the American accented story (Figure 7-B). In TRFs to the Dutch accented story, increased aptitude was associated with decreased sustained responses to acoustic features at relatively late lags (224–277 ms, *p*=.048; Figure 7-D), similar to the effect of proficiency described above (cf. Figure 6).

**Figure 7.**
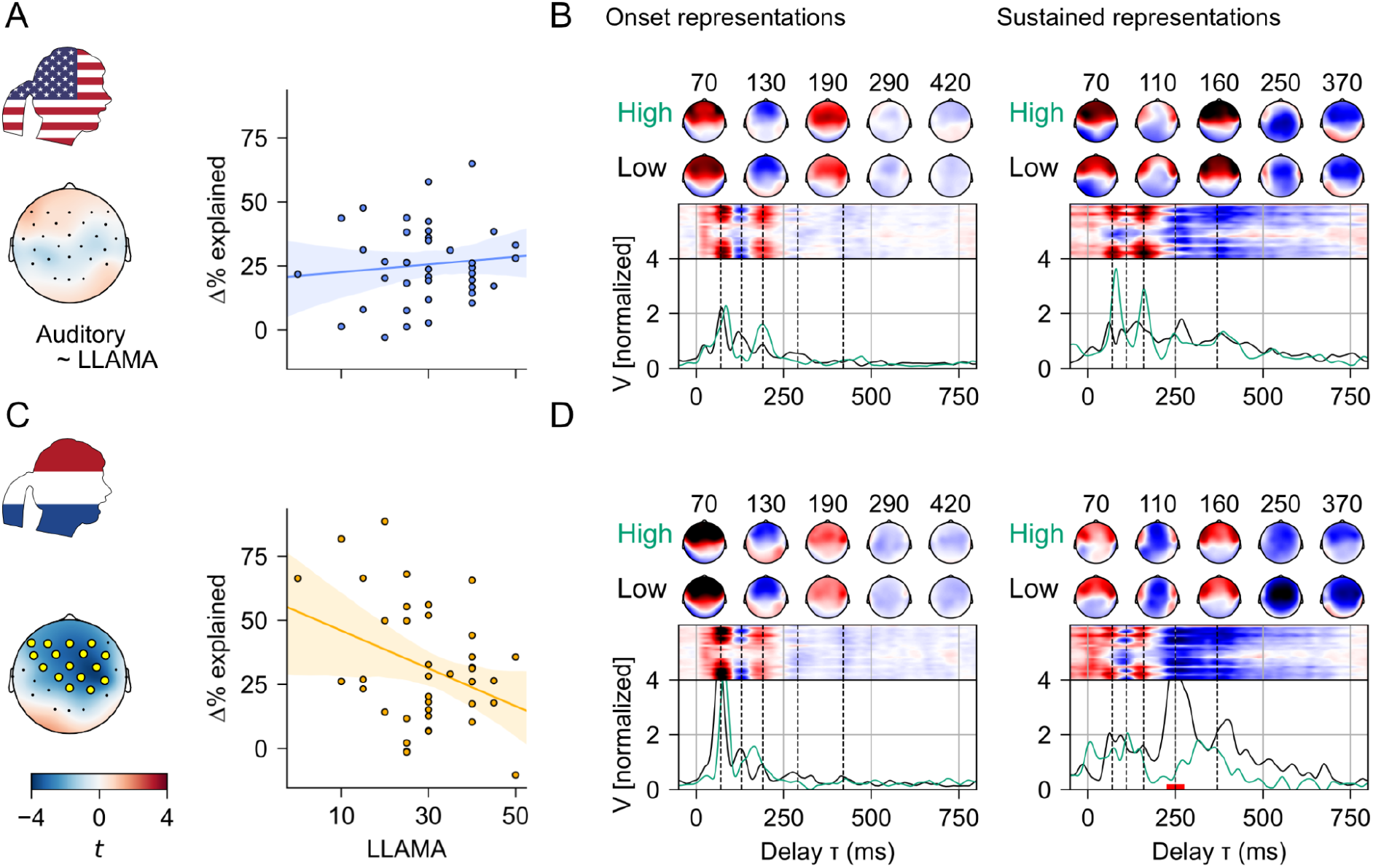
Auditory responses are modulated by acoustic-phonetic aptitude when listening to Dutch accented speech only. Unless mentioned otherwise, details are as in Figure 6. (A) Because predictive power at no sensor was meaningfully related to the LLAMA score (all *t*<2), the scatter-plot shows data for the average of all sensors. (B) Consistent with results from predictive power, TRFs were not significantly modulated by phonetic ability. Line plots show predictions for LLAMA_D scores of 10 and 50. (C) In brain responses to Dutch accented speech, increased phonetic ability was associated with smaller predictive power of auditory predictors, i.e., with weaker auditory responses. (D) TRFs related to sustained auditory representations of Dutch accented speech were modulated by aptitude at relatively late lags (224–277 ms).

Thus, Dutch listeners with higher acoustic-phonetic aptitude exhibited reduced acoustic responses when listening to English spoken with a Dutch accent. However, acoustic-phonetic aptitude did not affect acoustic responses when listening to English accented speech.

#### 3.4.3 English proficiency reduces sublexical representations of American-accented speech, and enhances sublexical representations of Dutch- accented speech

The predictive power of the English sublexical model (*sublexicalE*) was significantly associated with language proficiency, and this effect was modulated by speaker accent (Table 2). Proficiency affected responses in both American and Dutch accented speech (A: *χ*^2^(28)=49.62, *p*=.007; D: *χ*^2^(28)=63.81, *p<*.001). The interaction is illustrated in Figure 8. When listening to the American accented speaker, higher proficiency was associated with a decrease in predictive power, with large effects at frontal sensors bilaterally (Figure 8-A); in contrast, when listening to the Dutch accented speaker, proficiency was associated with an increase in predictive power primarily at right frontal sensors (Figure 8-C). Thus, when listening to English spoken with an American accent, more proficient listeners show less activation of English sublexical statistics compared to listeners with low proficiency; on the other hand, when listening to a Dutch accent, more proficient listeners activate English sublexical statistics more strongly.

**Figure 8.**
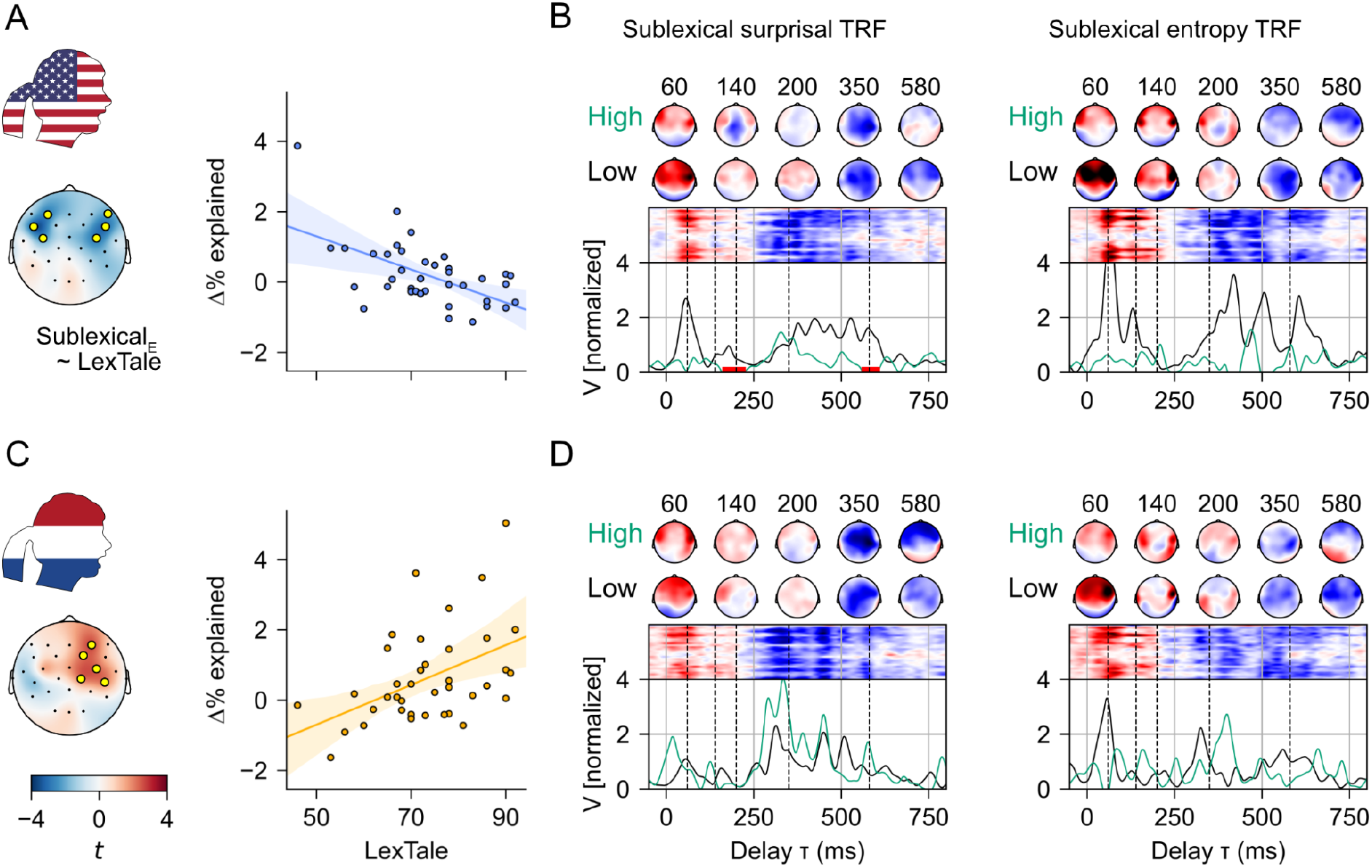
Activation of the English sublexical language model is modulated by proficiency and speaker accent. Unless mentioned otherwise, details are as in Figure 6. (A) For American accented English, higher proficiency is associated with reduced sublexical responses. (B) TRFs to the surprisal and entropy predictors based on the English sublexical language model. Surprisal is associated with a decreased response in more proficient listeners. (C) For Dutch accented English, higher proficiency is associated with stronger representation of the sublexical language model. (D) TRFs do not show significant effects of proficiency.

To determine *how* brain responses lead to this modulation of predictive power, we analyzed the corresponding TRFs, shown in Figure 8-BD. Here, a TRF reflects the component of the brain response to phonemes that scales with the corresponding predictor’s value, i.e., surprisal or entropy. The TRF to sublexical surprisal in the American accented story exhibit increased responses in listeners with low proficiency in middle (160–226 ms, *p*=.003) as well as later parts of the response (558–609 ms, *p*=.007). This suggests that the stronger activation of the sublexical model in individuals with low proficiency is due to increased extended cortical processing. On the other hand, the TRFs to the Dutch accented story do not exhibit a significant effect of LexTale, and thus do not provide a clear explanation for higher predictive power in high proficiency individuals.

#### 3.4.4 No evidence for a decrease in native language interference with increasing proficiency

Even though effects of native language interference persist in highly proficient non-native listeners (Garcia Lecumberri et al., 2010), we hypothesized that the magnitude of the interference might decrease with increasing proficiency. However, the predictive power of the models of native language interference (the *sublexicalD* predictor and the *word-form*ED>E contrast) were not significantly related to LexTale. Figure 9 shows plots of native language interference as a function of proficiency. The evidence for native language interference was averaged at 18 anterior sensors (manually selected, based on the observation that predictive power of the relevant comparisons was strongest in this region, cf. Figure 3). Even though some of the regression lines seem to exhibit a negative trend, none of these associations were significant (Table 2). This suggests that in the range of proficiency studied here, native language interference does not significantly decrease with increased proficiency.

**Figure 9.**
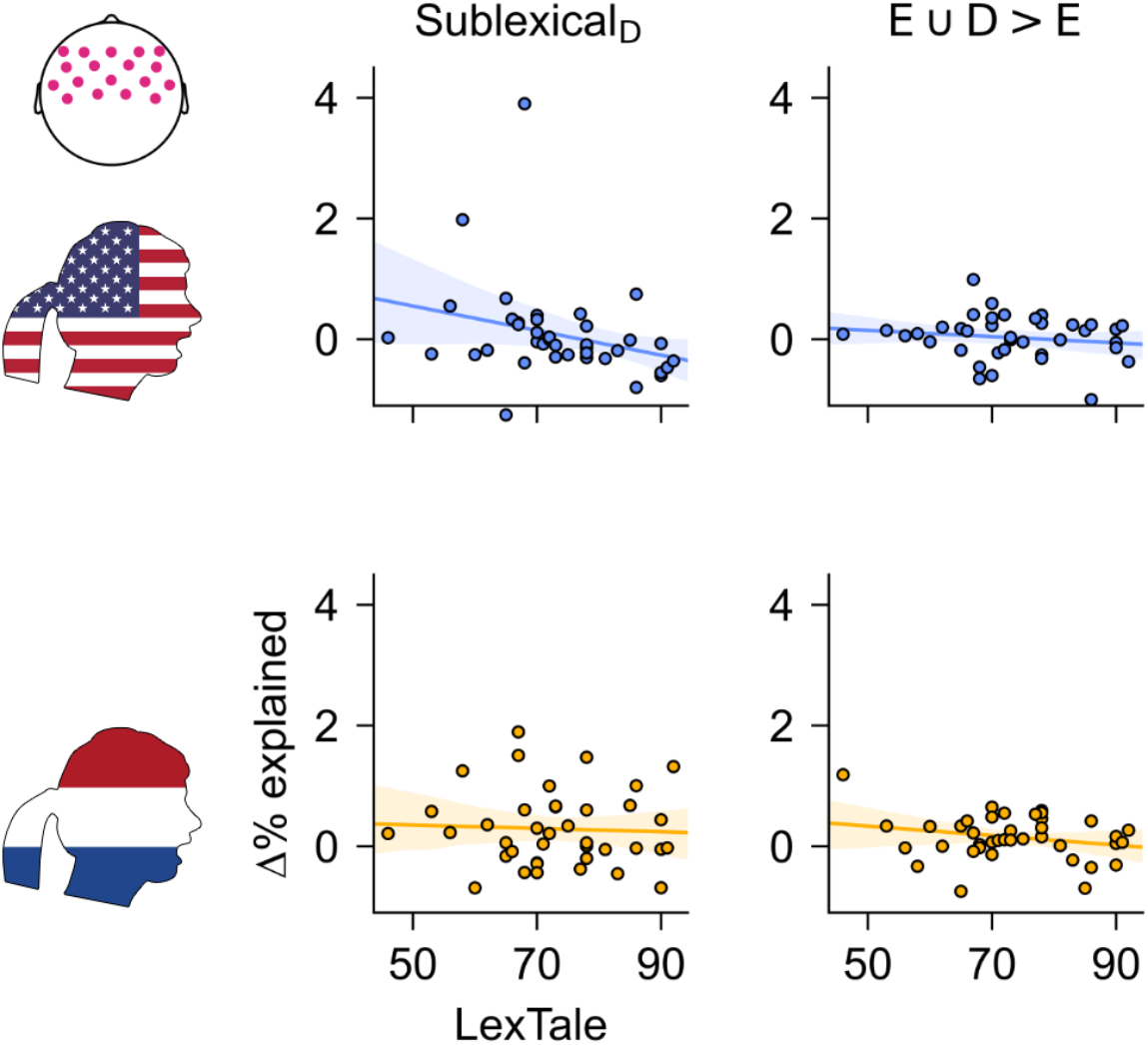
EEG responses that quantify the influence of the native language on non-native speech processing were not significantly related to proficiency. SublexicalD quantifies activation of the Dutch phoneme sequence model (i.e., comparison Equation 7 > Equation 6); E∪D>E quantifies the increase in predictive power due to including Dutch word forms (i.e., comparison Equation 9 > Equation 7). Data shown on the y-axis correspond to the average predictive power at anterior sensors (top left, pink sensors). Even though some regression plots seem to exhibit a negative trend, associations were not significant.

## 4 Discussion

### 4.1 Parallel predictive processing of non-native speech

When native speakers of English listen to their native language, they maintain predictive language models at different levels in parallel, including sublexical, word-form, and sentence models (Brodbeck et al., 2022; Z. Xie et al., 2023). We found that EEG responses of native speakers of Dutch, listening to an English story, exhibited evidence for parallel activation of similar language models. This suggests that parallel activation of different language models is also a property of non-native speech perception, at least at the intermediate and higher level of proficiency represented in our sample.

### 4.2 Activation of the native language

We found evidence for two ways in which the native language (Dutch) influenced brain responses associated with non-native (English) speech processing. First, listening to English activated a predictive model of Dutch phoneme sequences, in addition to the appropriate English phoneme sequence model. The interference effect was only significant in Dutch accented speech (although the evidence for a difference by speaker accent was weak). This suggests that listeners were not able to completely “turn off” statistical expectations based on phoneme sequence statistics in their native language, at least when listening to English spoken with a Dutch accent.

Second, brain responses to Dutch accented speech exhibited evidence for a word-form processing model that activates not only English words but Dutch and English words together. This effect provides a neural correlate for a phenomenon seen in behavioral results, showing activation of words from the native language during non-native listening (Hintz et al., 2022; Marian & Spivey, 2003; Spivey & Marian, 1999; Weber & Cutler, 2004). However, in our results this effect was significant only for Dutch accented speech, and was not detectable for English accented speech. Thus, in this more naturalistic listening scenario, the activation of words from the native language specifically depended on the accent. This may be because Dutch phonetic categories are inherently linked to Dutch lexical items more strongly than the newly learned American phonetics, or becuase the Dutch accent makes Dutch more salient in general and thus primes Dutch lexical competitors. We also surmise that, compared to naturalistic listening, the effect may be exaggerated in visual world studies, in which the native language competitors may be primed due to their presence on the visual display.

Neither of the effects of native language interference was modulated by proficiency. This suggests that this interference does not disappear in more proficient listeners. This neuroscientific evidence is consistent with previous behavioral results suggesting that native language interference persists even in advanced non-native listeners (Hintz et al., 2022; Spivey & Marian, 1999; Weber & Cutler, 2004). Together with our finding of increased native language interference in a non-native accent from one’s own language, this could explain why such a non-native accent becomes relatively more challenging at higher proficiency (Gordon-Salant et al., 2019; Pinet et al., 2011; X. Xie & Fowler, 2013): At lower proficiency, the non-native accent bestows an advantage due to the familiar acoustic-phonetic structure. At higher proficiency, the acoustic-phonetic structure of the native accent becomes more familiar, thus reducing the initial advantage of the non-native accent. Now, the disadvantage due to the persistently increased native language interference in the non-native accent becomes the dominant factor, making the non-native accent relatively more difficult than a native accent.

### 4.3 Acoustic representations are reduced by proficiency

More proficient listeners exhibited reduced amplitudes in brain responses to acoustic features. Our result replicates an earlier finding (Zinszer et al., 2022) and further suggests that this was primarily due to a reduction in late (>200 ms) responses. We broadly interpret this to indicate that in more proficient listeners, less neural work is being done with the acoustic signal at extended latencies. A potential explanation is that, when lower level signals can be explained from higher levels of representation, the bottom-up signals are inhibited (Rao & Ballard, 1999; Tezcan et al., 2022). Under these accounts, the observed result could reflect that more proficient listeners get better at explaining (and thus inhibiting) acoustic representations during speech listening. This would explain why the reduction is found primarily in late responses: Early responses reflect bottom-up processing of the auditory input and are similar across participants, but more proficient listeners have better acoustic-phonetic models that more quickly explain the bottom-up signal and thus inhibit the later responses.

### 4.4 Acoustic representations of Dutch accented English are reduced by acoustic-phonetic aptitude

Listeners that scored high on the LLAMA_D test of acoustic-phonetic aptitude also exhibited reduced auditory responses, but only in Dutch-accented English. As with proficiency, this affected primarily later response components (>220 ms). Similarly to the effect of proficiency, the reduced responses may indicate a reduction in neural work or better acoustic-phonetic models. The interaction with speaker accent, then, would indicate that acoustic-phonetic aptitude facilitates the recognition of English language words based on Dutch phonemes, and is less relevant for the American accent. This might sound counterintuitive, however, Dutch people tend to be exposed more to native English accents (e.g. through subtitled movies) than to Dutch accented English. Consequently, it might be that the Dutch accent is to some extent less naturally mapped to English word forms than the American accent.

### 4.5 Sublexical processing of the foreign language

Sublexical processing of English was modulated by proficiency in a complex manner, depending on the speaker’s accent: When listening to the story spoken with an American accent, increased proficiency was associated with *decreased* activation of the English sublexical language model. This is consistent with a previous report on Chinese non-native listeners, where increased English proficiency was associated with smaller responses related to a phonotactic measure (Di Liberto et al., 2021). Our results replicate this effect in Dutch non-native listeners, and tie it to sublexical (vs. word-form) processing. However, our results also suggest that the effect depends on the speaker’s accent: when listening to the story spoken with a Dutch accent, increased proficiency was associated with *increased* activation of the English sublexical model.

Interestingly, behavioral data indicate a similar interaction of proficiency with speaker accent: low proficiency listeners benefit from an accent corresponding to their own native language, whereas more proficient listeners benefit more from an accent native to the target language (Pinet et al., 2011; X. Xie & Fowler, 2013). Thus, as more proficient non-native listeners have tuned their phonetic perception more to a native accent (Di Liberto et al., 2021; Eger & Reinisch, 2019), phonetic cues in the non-native accented speech may become relatively less reliable. This may be due to the mismatch of the acoustic cues with the stored acoustic representations, but also due to the persistent native language interference (see above). This perceived reliability may influence the degree to which listeners rely on expectations from short-term transition probabilities between phonemes (i.e., the sublexical model) to provide a prior for interpreting the acoustic input: Decreased activation of the sublexical language model when listening to a native speaker might indicate that more proficient listeners rely less on this lower level model, perhaps also because the higher level models work more reliably and they don’t expect to need to fall back on the sublexical model as much. In contrast, the increase in activation of the sublexical language model when listening to the non-native accent may indicate that more proficient listeners increasingly recruit the sublexical language model to provide a prior for the imperfect bottom-up signal.

### 4.6 Lack of modulation of sentence level responses

We found no relationship between proficiency and responses related to the sentence-level language model. This finding suggests that listeners across our sample (intermediate to higher proficiency) used the English multi-word context to predict upcoming words and phonemes. This may indicate that listeners develop predictive models early on during non-native language learning (Frost et al., 2013; Sanders et al., 2002), especially when languages are structurally similar (Alemán Bañón & Martin, 2021). This may have been further facilitated by the language experience of our sample: English is frequently encountered in the Netherlands, which may have provided enough input even for intermediate proficiency listeners to develop models of language statistics.

### 4.7 Conclusions

We found relatively stable higher level neural language model activations (word-form and sentence level) from intermediate to high proficiency listeners, but reductions in the activation of auditory and sublexical representations with increased proficiency. This may indicate that listeners of intermediate proficiency are able to extract and use sentence level information appropriately in the non-native language (at least in the context of listening to the relatively easy story), but keep refining computations related to lower level acoustic and sublexical representations.

We also found evidence for a continued influence of native language statistics during naturalistic non-native listening. However, our results suggest a significant influence only in Dutch accented speech, where the Dutch speech sounds may increase activation of Dutch language representations. This selective interference may explain why a Dutch accent becomes relatively more challenging for highly proficient listeners. On the other hand, behavioral research may have inadvertently increased native language interference by increasing meta-linguistic awareness (Freeman et al., 2021), or by priming native language distractors.

## Acknowledgements

The data was collected as part of a VIDI grant from the Netherlands Organization for Scientific Research (NWO; grant number 276-89-003) awarded to O.S. C.B. was supported by NSF BCS- 1754284 and IIS-2207770, and a University of Connecticut seed grant from the Institute of the Brain and Cognitive Sciences.

